# Plant-plant interactions in wheat mixtures modulate mean and variance of susceptibility to Septoria tritici blotch

**DOI:** 10.1101/2024.08.26.609704

**Authors:** Laura Mathieu, Aurélie Ducasse, Elsa Ballini, Jean-Benoît Morel

## Abstract

Varietal mixtures are a promising agro-ecological approach to stabilizing yields by reducing diseases. The effects of mixtures stem from modifications of epidemiological processes and plant-plant interactions, which could explain some of the paradoxical observations made in the field. However, the role of plant-plant interactions in modifying bread wheat and durum wheat susceptibility to septoria tritici blotch remains to be elucidated. Our study aimed to determine the effect of such plant-plant interactions, by producing full matrices of binary mixtures in the absence of epidemics, on septoria symptoms—specifically necroses (lesions) and pycnidia (spore-containing structures). We employed statistical modeling to compare the mean and variance of focal plants’ phenotype in all mixtures *versus* pure conditions and in each mixture *versus* pure condition. Our findings demonstrate significant effects of plant-plant interactions on wheat susceptibility to septoria. Notably, these interactions had specific rather than general effects, with some but not all genotypic combinations significantly influencing focal susceptibility to septoria. Furthermore, mixtures resulted in reduced necrosis with lower variance, but increased pycnidia formation. These results reinforce the need to consider specific plant-plant interactions for their contribution to trait means and variances. Better considering these interactions could improve crop management strategies that enhance disease control.

**Highlight:** Wheat varietal mixtures modulate the mean and variance of septoria disease symptoms through specific plant-plant interactions. Mixtures reduce lesions and their variance but increase the formation of spore-containing bodies.

## Introduction

One promising approach to enhance the sustainability of agricultural systems is the adoption of varietal mixtures, which involves growing multiple cultivars within the same field. The renewed interest in varietal mixtures stems from their capacity to sustainably manage diseases (Mundt, 2002; Mundt *et al*., 2002). In wheat, this phenomenon has been observed across various devastating foliar diseases (Figueroa *et al*., 2018) such as leaf rust (Cox *et al*., 2004), yellow rust (Huang *et al*., 2012) and septoria (Kristoffersen *et al*., 2019). Varietal mixtures not only increase yield and decrease disease susceptibility but they also reduce yield variations (Borg *et al*., 2018). Indeed varietal mixtures have demonstrated potential in bolstering yield stability over time (Reiss and Drinkwater, 2018), and lessening environmental influences on crop productivity (Kiær *et al*., 2012; Moutier *et al*., 2022).

The positive outcomes of biodiversity on disease reduction are generally attributed to epidemiological shifts within mixtures (Borg *et al*., 2018). In particular, the spatial alternance of susceptible and resistant plants in mixtures generates a dilution effect that slows down epidemics. Some epidemiological parameters could also be directly modulated by the architecture produced in cultivar mixtures (Vidal *et al*., 2017). However, paradoxical effects of mixtures on disease susceptibility (Borg *et al*., 2018; Reiss and Drinkwater, 2018) are also observed. This was particularly observed in the case of the devastating Septoria tritici blotch (STB, caused by *Zymoseptoria tritici*) (Kristoffersen *et al*., 2022). While the majority of the mixtures (82%) exhibited a significant reduction of STB, the remaining 18% of the mixtures showed a 13% increase of susceptibility. Similarly, (Montazeaud *et al*., 2022) found that only 76% of mixtures were less affected by STB but that 24% were more diseased. These observed positive and negative effects in cultivar mixtures were attributed to underlying plant-plant interactions. Consistently, a wheat locus was identified such that its allelic diversity in the mixture was detrimental to STB resistance. The quantitative effect of this locus was as high than the observed, general effect of mixtures (17% disease reduction). These findings strongly support the idea that other processes than the epidemiological ones are also involved in regulating STB susceptibility in wheat varietal mixtures. Although plant-plant interactions have been hypothesized to modulate this pathosystem, only limited evidence supports this hypothesis since the effects of plant-plant interactions and epidemiological processes are often entangled in the field.

Moreover, the above-mentioned reduction of yield variance observed in mixtures across time (Reiss and Drinkwater, 2018) has been attributed to the buffering effects of ecological processes mobilized through diversity. Some reports also suggest that mere plant-plant interactions generated by plant diversity can reduce the average variance of individual traits (Siefert *et al*., 2015). In the same way, it was demonstrated that plant-plant interactions can change the average levels of disease susceptibility in varietal mixtures (Pélissier *et al*., 2023). Therefore, it is possible that plant-plant interactions could modify the variance of disease-related traits and thus contribute, to an integrative level, to a reduction of these trait variations along time or space.

The effects of plant-plant interactions are difficult to observe in the field where many ecological processes are undergoing. Experimental settings under controlled conditions are better suited for disentangling direct plant-plant interactions from other processes. Using greenhouse experiments and controlled infections excluding epidemiological processes, it was possible to demonstrate that direct plant-plant interactions are important modulators of disease susceptibility (Pélissier *et al*., 2021). For instance, in a specific case of varietal mixture of durum wheat, the presence of a non-inoculated intraspecific neighboring plant can increase susceptibility to STB (Pélissier *et al*., 2021). More recently, it was shown that plant-plant interactions can generally reduce the average susceptibility to leaf rust in durum wheat (Pélissier *et al*., 2023). Thus, general effects of plant-plant interactions on disease susceptibility exist. However, the general effects of plant-plant interactions on STB development are yet unknown despite the importance of this disease (Jørgensen *et al*., 2014).

The part of disease susceptibility phenotype in one given focal plant that is linked to a neighboring genotype can easily be assessed by decomposing the genotypic effects. Disease susceptibility can indeed be modeled by integrating direct genetic effects (DGE) and indirect genetic effects (IGE) (Wolf *et al*., 1998). DGE refers to the effect of the focal genotype on its own phenotype. In the case of a disease phenotype, DGE represents the genotype’s susceptibility (or conversely resistance). Currently, plant breeding focuses on DGE, with major resistance genes being key in controlling resistance. IGE is the general effect of the genotype of an adjacent plant on the susceptibility to so-called focal plants (i.e. those on which phenotype is considered). However, the neighbor genotype could also have a specific effect on the phenotype of only certain focal plants. This specific effect, which depends on its genotype and the genotype of the focal plant, is modeled by the DGE:IGE interaction. Identifying IGE could simplify the breeding programs, by selecting for general neighboring effects, while identifying DGE:IGE interaction would require more complex designs to test specific partner mixtures. Such model can be used to analyze the phenotypic data produced in experiments where all selected genotypes are grown in binary mixtures with all possible genotypes.

In this study, we analyzed the effect of plant-plant interactions on STB in binary mixtures of bread wheat on the one hand and durum wheat on the other. The aim was to investigate the type of interactions occurring in these mixtures and their effects on the mean of STB symptoms. Furthermore, to test the hypothesis that plant-plant interactions could contribute to the stability of traits in mixtures, we also considered the variance of STB symptoms. Two complete experimental matrices were produced, each containing all possible binary mixtures among 12 bread wheat varieties or 9 durum wheat varieties. For each matrix, two disease-associated phenotypes, necroses and pycnidia production, were measured. Statistical modeling was used to estimate DGE, IGE, and their interaction.

## Material and methods

### Plant material

The selected bread wheat and durum wheat genotypes analyzed in this study are varieties grown in France and were chosen for their susceptibility to the pathogen strains used. This criterion enabled us to study changes in basal immunity rather than resistance genes as in (Pélissier *et al*., 2023). A total of 12 bread wheat varieties and 9 durum wheat varieties were included in the analysis.

### Plant growth conditions

The wheat plants were cultivated in a greenhouse at PHIM (Montpellier, France) under controlled conditions. The photoperiod was 16 hours of light and 8 hours of darkness, with a temperature of 24°C/20°C and a light intensity of 250 μmol/s/m2. The plants were grown in pots measuring 8cm x 8cm x 8cm, filled with a mixture of 50% vegetable soil, 50% Neuhaus N2 soil, and 133.3g TopPhos per 100% soil.

### Experimental design

The experiment was conducted between October and December 2020 for durum wheat and between November 2021 and February 2022 for bread wheat. Two complete matrices of binary mixtures were produced from 12 varieties for bread wheat and 9 varieties for durum wheat. All genotypic combinations, including mixtures and pure conditions, were grown. Therefore, there were 144 combinations (among which were 12 pure conditions) for bread wheat and 81 for durum wheat (among which were 9 pure conditions). Each combination was repeated at least four times. Both genotypes were phenotyped in each binary mixture, alternating between being the focal plants, on which symptoms are observed, and the neighboring plants symmetrically. Each pot contained two lines of four plants, each corresponding to one genotype. The pots were randomly placed in the greenhouse.

### Inoculation and symptom assessment

Bread wheat plants were inoculated with the STB strain IPO9415, while durum wheat plants were inoculated with the P1A strain (Ballini *et al*., 2020). For both, an inoculum of 10^6^ spores per mL was prepared and applied to the last ligated leaf 21 days after sowing using a brush. The plants were then placed in transparent plastic bags for three days to saturate the air humidity and promote the pathogen’s growth. STB symptoms were assessed 17 days after inoculation using an Epson Perfection V370 Photo scanner. Two phenotypes of interest were analyzed: the surface area of necrosis, which refers to the lesions, and the necrosis area containing pycnidia, which are the structures containing spores. The symptoms were scored using SeptoSympto for bread wheat (Mathieu *et al*., 2024) or manually for durum wheat due to the lower resolution of images. This was done to estimate the necrosis area and the necrosis area containing pycnidia.

### Statistical analyses

#### Model for estimating DGE, IGE and their interaction

The direct genetic effect (DGE, i.e. effect of focal identity), the indirect genetic effect (IGE, i.e. effect of neighbor identity) and their interaction (DGE:IGE, i.e. interaction between focal identity and neighbor identity) were estimated with a non-parametric linear model, since the necrosis and pycnidia phenotypes do not follow the assumptions of error normality and homoscedasticity. An aligned and ranked linear model (Equation 1), using the ARTool package (Kay and Wobbrock, 2016), allowed these different effects to be estimated with an Aligned Ranked Transform (ART) ANOVA.

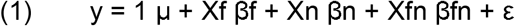

where y is the vector of phenotypic observations, Xf, Xn and Xfn are known incidence matrices and μ is the mean phenotype. βf, βn and βfn correspond to fixed effects for focal genotype, neighbor genotype and interaction between focal genotype and neighbor genotype, respectively.

### Comparison of focal susceptibility between each mixture and the pure condition

The susceptibility of each focal genotype to septoria was compared between each mixture and the pure/monoculture condition. The mean ratio was calculated as xij / xii’, where xij is the adjusted mean of focal genotype i grown with neighboring genotype j and xii’ is the adjusted mean of focal genotype i grown with the same neighboring genotype i’ (here i’ means j=i, as this is a pure condition). Adjusted means are least squares means calculated from the aligned and ranked linear model (M2) using the emmeans package (Lenth, 2022). The variance ratio was calculated as vij / vii’, where vij is the variance of focal genotype i grown with neighboring genotype j and vii’ is the variance of focal genotype i grown with the same neighboring genotype i’ (pure condition). Means were compared using contrast tests following an aligned and ranked linear model (Equation 2), while variances were compared using Flingner tests. To correct for multiple testing, a Bonferroni correction was applied for each focal genotype.

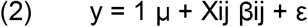

where y is the vector of phenotypic observations, μ is the mean phenotype and Xij is the known incidence matrix. βij corresponds to the fixed effect for the genotypic combination of focal i and neighbor j.

### Comparison between all mixtures and pure conditions

Septoria susceptibility was compared between mixtures and pure conditions using a Wilcoxon test for medians and a Fligner test for variances.

## Results

### Susceptibility to STB is modulated in wheat binary mixtures in the absence of epidemics

Wheat susceptibility to septoria was first considered between mixtures and pure conditions as a whole in order to test for global effects of mixtures. Bread wheat mixtures showed significantly fewer necroses than pure conditions, despite a relatively small difference (12% less necrotic area in mixtures; Fig. 1A). Durum wheat mixtures did not differ significantly from pure conditions (Fig. 1C).

**Fig. 1.**
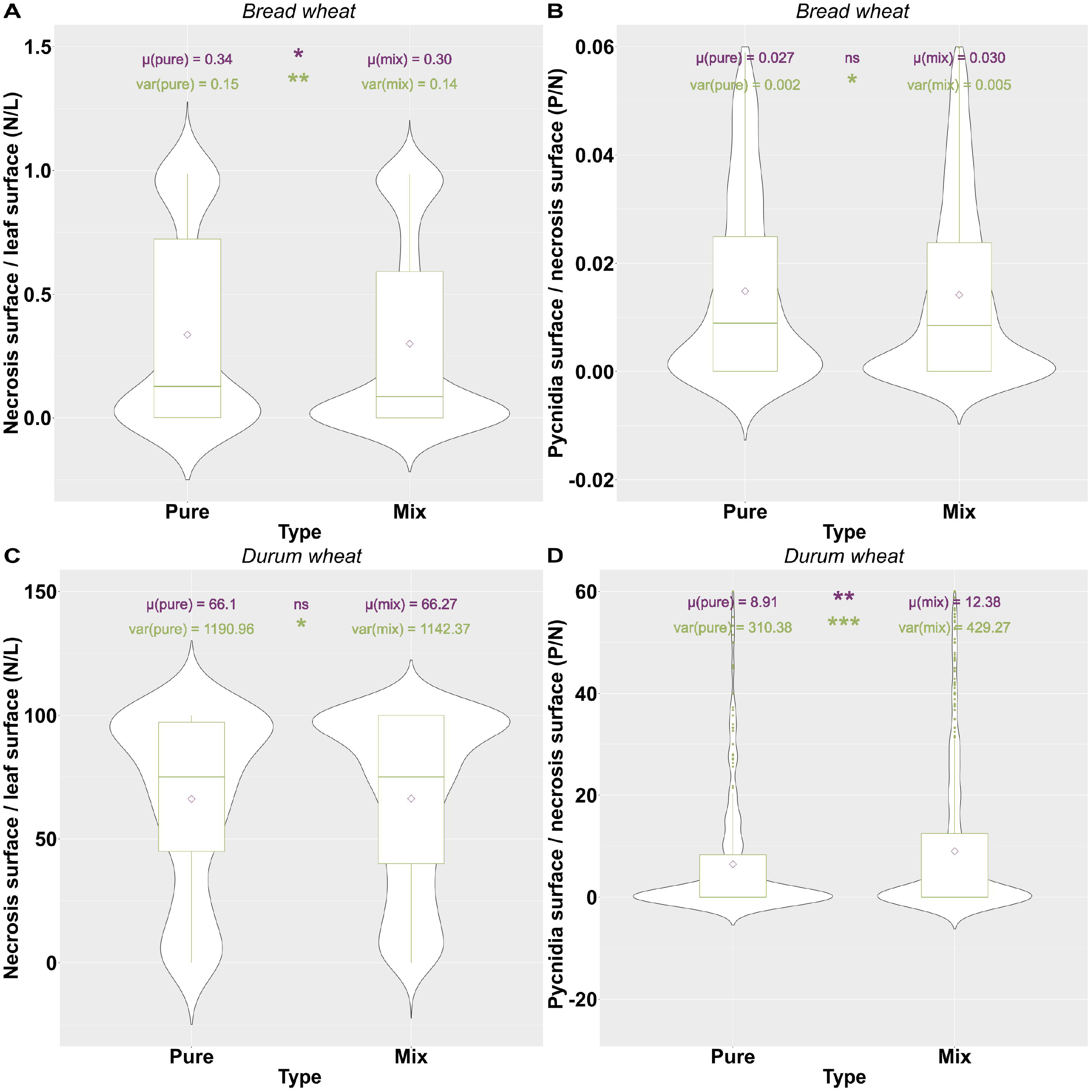
Wheat susceptibility to Septoria tritici blotch (STB) in pure conditions and mixtures. **A** Necrosis phenotype in bread wheat. **B** Pycnidia phenotype in bread wheat. **C** Necrosis phenotype in durum wheat. **D** Pycnidia phenotype in durum wheat. Statistical comparisons between wheat susceptibility in all mixtures (mix) and pure conditions (pure) were conducted using Wilcoxon tests for medians and Fligner tests for variances. Significance levels are denoted as follows:. : p < 0.1, *: p < 0.05, **: p < 0.01 and ***: p < 0.001.

Mixtures produced more pycnidia than pure conditions for both species (Fig. 1B-D). Specifically, there was a significant increase of 11% and 39% in pycnidia surface for respectively bread and durum wheat grown in mixtures compared to pure conditions. While pycnidia increased, necrosis areas were reduced in mixtures.

We then considered susceptibility to STB (necrosis and pycnidia) for focal genotypes grown with different genotypes as neighbors and compared to focal genotype grown with itself (pure controls) (Fig. 2). In the bread wheat matrix, 9% of the focal plants in mixtures exhibited a significantly different necrosis surface compared to the pure control (Fig. 2A). Among these cases, focal plants were less susceptible in 7 cases, while 5 cases showed increased susceptibility of focal plants in the mixture compared to the pure condition. At best, necrosis was significantly reduced by 100% in Fructidor and increased by 30% in Titlis when grown with Centurion. Pycnidia, on the other hand, were less affected by plant-plant interactions in bread wheat mixtures, with only a few mixtures exhibiting significant differences (Fig. 2B). Notably, when grown with the neighboring genotype Titlis, the focal genotype Centurion produced 87% less pycnidia, whereas the focal genotype Descartes produced 72% more pycnidia compared to pure conditions.

**Fig. 2.**
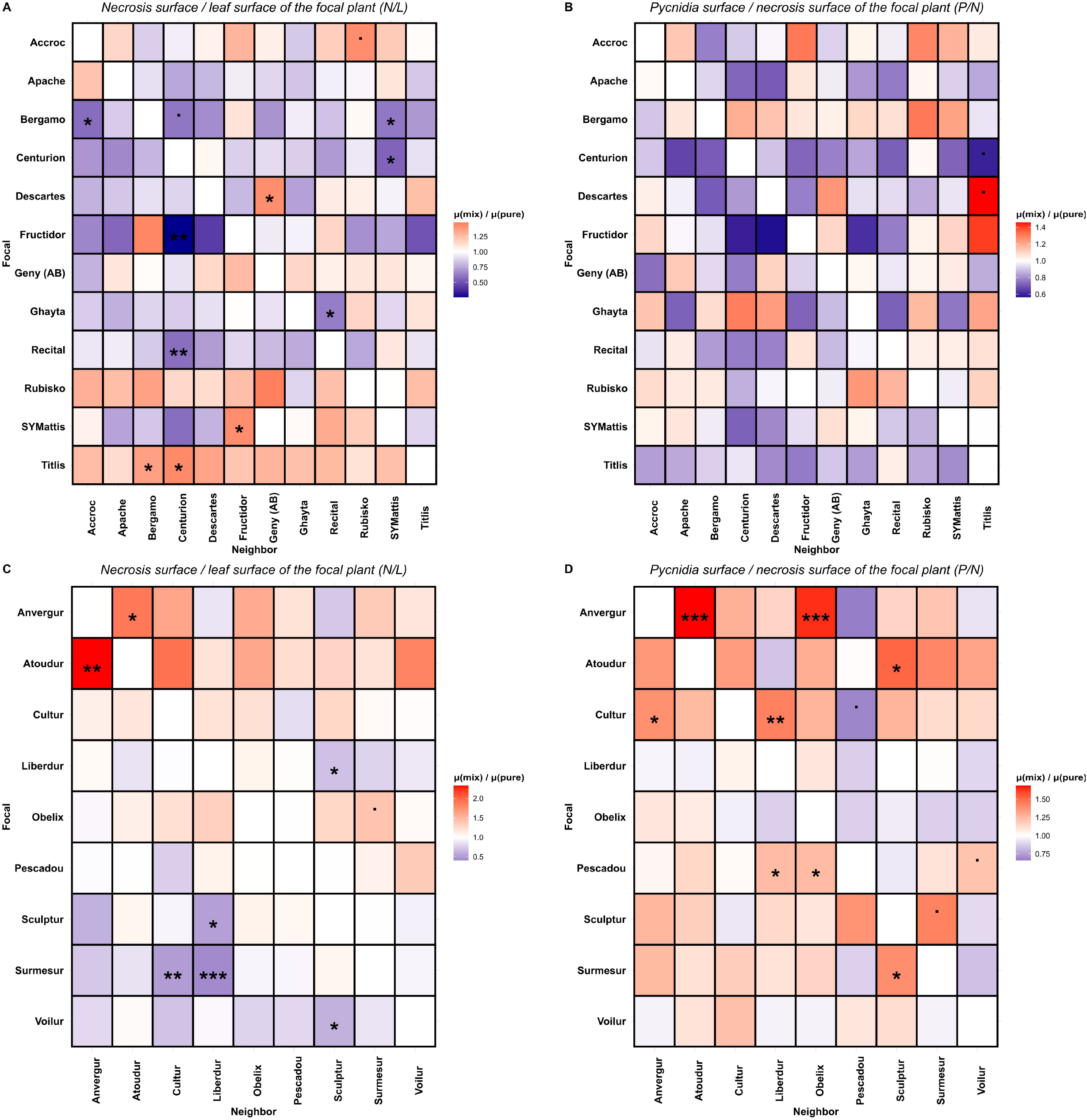
Modulation of focal average susceptibility to Septoria tritici blotch (STB) by neighboring plants. **A** Necrosis phenotype in bread wheat matrix. **B** Pycnidia phenotype in bread wheat matrix. **C** Necrosis phenotype in durum wheat matrix. **D** Pycnidia phenotype in durum wheat matrix. The ratio of focal susceptibility in each mixture compared to pure condition was calculated and depicted in the Fig; thus diagonal values are equal to 1. Cases where the focal susceptibility in mixture is higher than pure condition are indicated in red, while those with lower susceptibility are shown in blue. Statistical comparisons between focal susceptibility in each mixture and pure condition were conducted using non-parametric contrast tests. Significance levels are denoted as follows:. : p < 0.1, *: p < 0.05, **: p < 0.01 and ***: p < 0.001.

Similarly, in the durum wheat matrix, the necrosis area of focal plants was modified in ∼10% of mixtures compared to pure conditions (Fig. 2C). Among these cases, focal plants were less susceptible in 5 cases and more susceptible in 3 cases. The previously observed enhanced susceptibility of Cultur in the presence of Atoudur (Pélissier *et al*., 2021) was again detected. At best, necrosis was reduced by 56% in Surmesur grown with Liberdur and increased by 136% in Atoudur grown with Anvergur. The impact of plant-plant interactions on STB reproduction was more significant in durum wheat mixtures than in bread wheat, with 13.6% of mixtures showing differences in the necrosis area containing pycnidia of focal plants compared to pure conditions (Fig. 2D). Among these cases, only one showed a 57% significant decrease in pycnidia for the focal plants grown in mixtures. The Cultur focal plant produced 115% more pycnidia when grown with Liberdur and conversely less when grown with Pescadou compared to pure condition.

Therefore, for both bread and durum wheat, the results indicate a strong specific effect of the neighboring genotype on the focal susceptibility, which depends on the combination between its genotype and the focal genotype.

### Wheat susceptibility to STB in mixtures is mainly modified by specific plant-plant interactions

It was clear from these results that besides direct genetic effects (DGE; here an indirect measure of basal immunity) on susceptibility, mixtures also displayed general (Fig. 1) and specific (genotype-by-genotype) (Fig. 2) effects. We thus used a DGE:IGE model to quantify the respective importance of these effects on susceptibility. Table 1 provides a summary of the proportion of phenotypic variance in focal plants explained by these effects and their statistical significance. DGE was found to be significant across all matrices, explaining a substantial proportion of the phenotypic variance (ranging from 14.6% to 40%). Plant-plant interactions were estimated using the IGE (general neighbor effect) and the DGE:IGE interaction (specific effect). While IGE was significant for both phenotypes in bread wheat and durum wheat matrices, it explained a relatively small proportion of phenotypic variance (ranging from 1.3% to 5.7%). In contrast, the DGE:IGE interaction accounted for a substantial proportion of phenotypic variance, ranging from 9.6% to 17.4%. Notably, this interaction can even exceed the influence of DGE alone in the necrosis phenotype of the bread wheat matrix. These results demonstrate that while there may be a general neighbor effect (IGE), the modulation of susceptibility to septoria in focal plants depends on the precise combination of mixed genotypes, indicating the predominance of specific plant-plant interactions.

**Table 1:** Modulation of wheat susceptibility to Septoria tritici blotch (STB) by genetic effects. Direct genetic effects (DGE) refer to the resistance of focal plants, while indirect genetic effects (IGE) and DGE : IGE interaction encompass general and specific neighbor effects, respectively. Non-parametric linear models were used to estimate the percent of phenotypic variance explained by genetic effects and their significance. Significance levels are denoted as follows: *: p < 0.05, **: p < 0.01 and ***: p < 0.001.

| Population | DGE<br>Focal identity | IGE<br>Neighbor identity | DGE:IGE<br>Focal : neighbor interaction |
| --- | --- | --- | --- |
| Bread wheat - *Z. tritici*<br>Necrosis | 14,6%<br>*** | 4,3%<br>*** | 17,4%<br>*** |
| Bread wheat - *Z. tritici*<br>Pycnidia | 19,5%<br>*** | 3,7%<br>*** | 9,6%<br>*** |
| Durum wheat - *Z. tritici*<br>Necrosis | 17,6%<br>*** | 1,3%<br>* | 10,5%<br>*** |
| Durum wheat - *Z. tritici*<br>Pycnidia | 40,0%<br>*** | 5,7%<br>*** | 11,9%<br>*** |

### Mixtures show small but significant effects on variance

Besides affecting mean susceptibility, we also tested whether mixtures could modulate phenotypic variance. When mixtures and pure conditions were considered as a whole (Fig. 1), we could not test this hypothesis for pycnidia since the variances and means were positively related for this phenotype, a situation that does not allow testing significance of variance differences (i.e. the higher the mean is, the higher the variance is; Fig. S1). For necrosis, the means and variances were not related, allowing the testing of our hypothesis. The test revealed that for both bread wheat and durum wheat, the variance in mixtures was lower than in pure conditions, with 6,7% and 4% less variance respectively (Fig. 3A-C). These results highlight a slight but significant general effect of plant-plant interactions on the variance of plant susceptibility to STB in mixtures.

**Fig. 3.**
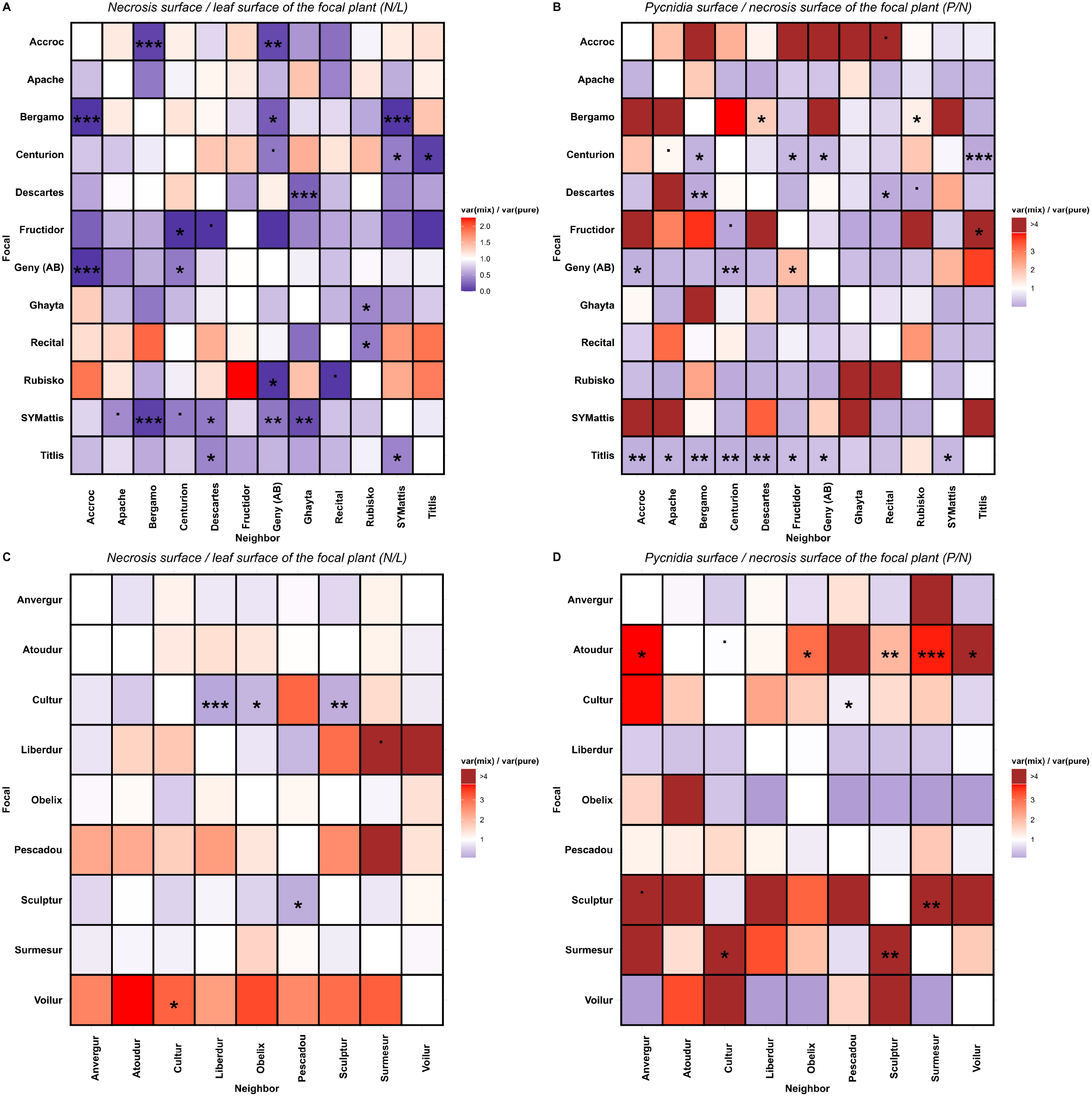
Modulation of focal susceptibility variance to Septoria tritici blotch (STB) by neighboring plants. **A** Necrosis phenotype in bread wheat matrix. **B** Pycnidia phenotype in bread wheat matrix. **C** Necrosis phenotype in durum wheat matrix. **D** Pycnidia phenotype in durum wheat matrix. The ratio of focal susceptibility variance in each mixture compared to pure condition was calculated and depicted in the Fig; thus diagonal values are equal to 1. Cases where the focal susceptibility variance in mixture is higher than pure condition are indicated in red, while those with lower susceptibility variance are shown in blue. Statistical comparisons between focal susceptibility in each mixture and pure condition were conducted using non-parametric Fligner tests. Significance levels are denoted as follows:. : p < 0.1, *: p < 0.05, **: p < 0.01 and ***: p < 0.001.

We then look, when mixtures and pure were considered through specific focal-neighbor genotype combinations (Fig. 3) and no longer as a whole (Fig. 1), if we could observe significant changes in variance between mixtures and pure conditions. Differences between variances of focal plant phenotypes in mixtures and pure conditions were obtained in the four data matrices (Fig. 3). In bread wheat, plant-plant interactions have a greater impact on variances than on means, with ∼19% of mixtures displaying significant differences in focal plant phenotype variance compared to pure conditions, all of which showing a decrease in variance (Fig. 3A). A similar trend was observed for pycnidia, with 17% of mixtures exhibiting modified variance of focal plant phenotype (Fig. 3B). The pycnidia variance of focal plants was significantly lower in 18 mixtures, and higher in 6 cases compared to the pure condition. In contrast, the durum wheat matrix appeared less affected by differences in variance due to plant-plant interactions than the bread wheat matrix, with 7% of mixtures showing significant differences in necrosis variance (Fig. 3C) and ∼14% showing differences in pycnidia variance (Fig. 3D). Specifically, there were 4 cases where focal plant necrosis was less variable than in pure condition and 2 cases where the opposite was observed. Additionally, out of 11 cases, only 2 showed a decrease in pycnidia variance of focal plants within mixtures. These variance matrices demonstrate that plant-plant interactions not only impact average phenotype but also phenotype variance. Just like for means, variance differences were also dependent on genotypic combinations.

## Discussion

### Plant-plant interactions modulate disease-related traits in wheat infected by STB

The present study sheds light on the direct impact of plant-plant interactions on wheat susceptibility to STB. Indeed, in our experimental set-up, epidemics are not possible and other ecological processes such as competition for light are very limited. A total of 33 (15,7%) of the 204 binary mixtures tested were affected for either necrosis or pycnidia production. This is similar to previous observations in durum wheat and rice (Pélissier *et al*., 2023). The findings reveal that these interactions have a strong specific (up to 17.4% in bread wheat and 11.9% in durum wheat), and a rather small general effect (up to 4.3% in bread wheat and 5.7% in durum wheat) on modulating wheat susceptibility to STB. Consistently, IGE was low and DGE:IGE was strong. These results are also similar to those found in the case of durum wheat and leaf rust (Pélissier *et al*., 2023) and reinforce the yet scarce reports demonstrating that neighbors can modulate susceptibility in their plant neighborhood. Thus, just like microbiome is now considered as a major player for modulating plant’s resistance to pathogens (Vannier *et al*., 2019), plant neighbors should be considered when designing crops with enhanced resistance. What remains to be determined is whether neighbor plants can modulate the effects of wheat resistance genes against STB. More generally, testing the effects of mixtures on major disease resistance genes will be important in the future to evaluate cultivar mixtures on these key genes for crop protection.

### Complex trait-trait combinations dominate the modulation of susceptibility to STB

Our observation of a small but significant IGE suggests that a constitutive mechanism for all neighbors may exist. However, we did not detect obvious cases of neighbors generally modulating all focal genotypes. Conversely, with the notable exception of Titlis, we did not identify a focal genotype that was generally affected by most neighbors. Our findings suggest that the mechanisms primarily influencing susceptibility to STB vary depending on the genotypic composition and, consequently, the traits of the mixed varieties. Our results differ from previous studies indicating that such specific interactions are not the main factors explaining yield-related phenotypes (Forst *et al*., 2019; Haug *et al*., 2023). These traits are yet to be identified but several speculative models can be proposed. One of these postulates that competition, and the associated access to resources, could modulate resistance in mixtures; for instance, a neighbor with large roots could outcompete the focal plant that would change its growth and consequently immunity since both are known to be related (de Tombeur *et al*., 2023). The observed phenotypic plasticity in STB susceptibility may be driven by different traits of the neighbors varying according to the genotypic combination and thus by resulting environmental changes. The detailed analysis and description of specific mixtures will help better understand the molecular mechanisms involved in the changes of susceptibility in mixtures.

### Plant-plant interactions and paradoxical effects of mixtures in the field

Comparisons of focal susceptibility in each mixture with the corresponding pure condition demonstrate that changes in susceptibility to STB can increase important parameters of epidemics. In particular, our study indicates that pycnidia production is increased by plant-plant interactions, as we observed a general positive effect of mixture on pycnidia production in durum wheat. In bread wheat, a similar but not significant effect was found. Pycnidia are structures containing spores responsible for dispersing the fungus and causing the epidemic (Suffert *et al*., 2011). With increasing pycnidia production, some plant-plant interactions could thus produce antagonistic effects to otherwise beneficial, ecological processes generated by cultivar mixtures. This may explain the variation in observed mixture effects in the field (Kristoffersen *et al*., 2022). This is also consistent with the finding that the diversity at a single wheat locus can increase disease (Montazeaud *et al*., 2022). Testing, in the field, the negative effects of some plant-plant interactions observed under controlled conditions, could confirm that such interactions are responsible for some of the unwanted, paradoxical observations made with cultivar mixtures.

### Could plant-plant interactions contribute to the observed stability of mixtures?

Phenotypic plasticity is defined as a change in the phenotype expressed by a single genotype in different environments. It has been mostly measured under changing, abiotic environments (Gratani, 2014) and not often measured in changing intra-specific neighborhoods (Dahlin *et al*., 2020). For instance, biomass, grain production or height were shown to be overall modified in barley mixtures. Similar to our findings, these authors also observed a strong neighboring effect. Furthermore, we could measure that not only the mean of susceptibility trait but also its variance was affected by plant-plant interactions. Indeed, necrosis variance was reduced in wheat mixtures. Such a change in trait variance within plant communities was already observed (Siefert *et al*., 2015). For instance, intraspecific trait variation has been found to decrease with increasing species richness for plant height or leaf traits (Siefert *et al*., 2015). Similarly, adaptative similarity (the fact that the values of a trait converge more in changing environments than on average) was observed in barley mixtures for various growth traits (Dahlin *et al*., 2020). However, no study to date has shown a reduction in the variance of disease susceptibility in plant mixtures. Our finding that necrosis variance is reduced in mixtures is reminiscent to observations made in communities of social insects. Indeed, variation in the immune response decreases with increased genetic diversity in bees (Simone-Finstrom *et al*., 2016). This decrease in disease variation suggests the presence of social behavior in response to diversity. Whether this reduction of trait variance contributes to the global, observed yield stability is still speculative; it raises the possibility that plant-plant interactions could somehow force traits to stay around the mean and thus contribute to trait stability in cultivar mixtures.

In summary, our study demonstrates that plant-plant interactions have a significant impact on wheat susceptibility to STB, affecting both the average susceptibility and variance. These findings highlight the importance of considering specific interactions in mixture design to improve crop management strategies and efficiently mitigate disease.

## Supplementary data

The following supplementary data are available at JXB online.

Fig. S1. Relation between average susceptibility and variance of focal plants.

## Acknowledgments

We are grateful to Alessandro Ramos for its technical help. We thank Jacques David and Bénédicte Fontez for their kind help with the statistical analyses.

## Author contributions

LM, EB and JBM conceptualized the study and designed the experiment. LM and AD performed the experiments. LM analyzed the data and led the writing of the manuscript. LM, JBM and EB reviewed the manuscript. JBM and EB supervised the work. All authors have read and approved the final manuscript.

## Conflict of interest

The authors declare that they have no competing interests.

## Funding

This work was supported by the French National Research Agency under the Investments for the Future Program [grant ANR-20-PCPA-0006, ANR-19-CE20-0005].

## Data availability

The phenotypic data underlying this article are available in GitHub at https://github.com/LauraC-Mathieu/wheat-STB-matrices.

## Figure legends

**Supplementary Fig. S1. Relation between average susceptibility and variance of focal plants. A** Necrosis phenotype in bread wheat. **B** Pycnidia phenotype in bread wheat. **C** Necrosis phenotype in durum wheat. **D** Pycnidia phenotype in durum wheat.

## Notes

### Competing Interest Statement

The authors have declared no competing interest.

## References

Ballini E, Tavaud M, Ducasse A, et al. 2020. Genome wide association mapping for resistance to multiple fungal pathogens in a panel issued from a broad composite cross-population of tetraploid wheat Triticum turgidum. Euphytica 216, 92.

Borg J, Kiær LP, Lecarpentier C, Goldringer I, Gauffreteau A, Saint-Jean S, Barot S, Enjalbert J. 2018. Unfolding the potential of wheat cultivar mixtures: A meta-analysis perspective and identification of knowledge gaps. Field Crops Research 221, 298–313.

Cox CM, Garrett KA, Bowden RL, Fritz AK, Dendy SP, Heer WF. 2004. Cultivar Mixtures for the Simultaneous Management of Multiple Diseases: Tan Spot and Leaf Rust of Wheat. Phytopathology® 94, 961–969.

Dahlin I, Kiær LP, Bergkvist G, Weih M, Ninkovic V. 2020. Plasticity of barley in response to plant neighbors in cultivar mixtures. Plant and Soil 447, 537–551.

Figueroa M, Hammond-Kosack KE, Solomon PS. 2018. A review of wheat diseases—a field perspective. Molecular Plant Pathology 19, 1523–1536.

Forst E, Enjalbert J, Allard V, Ambroise C, Krissaane I, Mary-Huard T, Robin S, Goldringer I. 2019. A generalized statistical framework to assess mixing ability from incomplete mixing designs using binary or higher order variety mixtures and application to wheat. Field Crops Research 242, 107571.

Gratani L. 2014. Plant Phenotypic Plasticity in Response to Environmental Factors. Advances in Botany 2014, 208747.

Haug B, Messmer MM, Enjalbert J, Goldringer I, Flutre T, Mary-Huard T, Hohmann P. 2023. New insights towards breeding for mixed cropping of spring pea and barley to increase yield and yield stability. Field Crops Research 297, 108923.

Huang C, Sun Z, Wang H, Luo Y, Ma Z. 2012. Effects of wheat cultivar mixtures on stripe rust: A meta-analysis on field trials. Crop Protection 33, 52–58.

Jørgensen LN, Hovmøller MS, Hansen JG, et al. 2014. IPM Strategies and Their Dilemmas Including an Introduction to http://www.eurowheat.org. Journal of Integrative Agriculture 13, p265–281.

Kay M, Wobbrock J. 2016. ARTool: aligned rank transform for nonparametric factorial ANOVAs. R package version 0.10 2.

Kiær LP, Skovgaard IM, Østergård H. 2012. Effects of inter-varietal diversity, biotic stresses and environmental productivity on grain yield of spring barley variety mixtures. Euphytica 185, 123–138.

Kristoffersen R, Eriksen LB, Nielsen GC, Jørgensen JR, Jørgensen LN. 2022. Management of Septoria Tritici Blotch Using Cultivar Mixtures. Plant Disease 106, 1341–1349.

Kristoffersen R, Jørgensen LN, Eriksen LB, Nielsen GC, Kiær LP. 2019. Control of Septoria tritici blotch by winter wheat cultivar mixtures: Meta-analysis of 19 years of cultivar trials. Plant Biology.

Mathieu L, Reder M, Siah A, Ducasse A, Langlands-Perry C, Marcel TC, Morel J-B, Saintenac C, Ballini E. 2024. SeptoSympto: a precise image analysis of Septoria tritici blotch disease symptoms using deep learning methods on scanned images. Plant Methods 20, 18.

Montazeaud G, Flutre T, Ballini E, et al. 2022. From cultivar mixtures to allelic mixtures: opposite effects of allelic richness between genotypes and genotype richness in wheat. New Phytologist 233, 2573–2584.

Moutier N, Baranger A, Fall S, Hanocq E, Marget P, Floriot M, Gauffreteau A. 2022. Mixing Ability of Intercropped Wheat Varieties: Stability Across Environments and Tester Legume Species. Frontiers in Plant Science 13.

Mundt CC. 2002. USE OF MULTILINE CULTIVARS AND CULTIVAR MIXTURES FOR DISEASE MANAGEMENT. Annual Review of Phytopathology 40, 381–410.

Mundt CC, Cowger C, Garrett KA. 2002. Relevance of integrated disease management to resistance durability. Euphytica 124, 245–252.

Pélissier R, Ballini E, Temple C, et al. 2023. The genetic identity of neighboring plants in intraspecific mixtures modulates disease susceptibility of both wheat and rice. (CH Haney, Ed.). PLOS Biology 21, e3002287.

Pélissier R, Buendia L, Brousse A, Temple C, Ballini E, Fort F, Violle C, Morel J-B. 2021. Plant neighbour-modulated susceptibility to pathogens in intraspecific mixtures. (R Napier, Ed.). Journal of Experimental Botany 72, 6570–6580.

Reiss ER, Drinkwater LE. 2018. Cultivar mixtures: a meta-analysis of the effect of intraspecific diversity on crop yield. Ecological Applications 28, 62–77.

Siefert A, Violle C, Chalmandrier L, et al. 2015. A global metaLanalysis of the relative extent of intraspecific trait variation in plant communities. (J Chase, Ed.). Ecology Letters 18, 1406–1419.

Simone-Finstrom M, Walz M, Tarpy DR. 2016. Genetic diversity confers colony-level benefits due to individual immunity. Biology Letters 12, 20151007.

Suffert F, Sache I, Lannou C. 2011. Early stages of septoria tritici blotch epidemics of winter wheat: buildLup, overseasoning, and release of primary inoculum. Plant Pathology 60, 166–177.

de Tombeur F, Pélissier R, Shihan A, et al. 2023. Growth-defence trade-off in rice: fast-growing and acquisitive genotypes have lower expression of genes involved in immunity. Journal of Experimental Botany, erad071.

Vannier N, Agler M, Hacquard S. 2019. Microbiota-mediated disease resistance in plants. PLOS Pathogens 15, e1007740.

Vidal T, Boixel A-L, Durand B, de Vallavieille-Pope C, Huber L, Saint-Jean S. 2017. Reduction of fungal disease spread in cultivar mixtures: Impact of canopy architecture on rain-splash dispersal and on crop microclimate. Agricultural and Forest Meteorology 246, 154–161.

Wolf JB, Brodie III ED, Cheverud JM, Moore AJ, Wade MJ. 1998. Evolutionary consequences of indirect genetic effects. Trends in Ecology & Evolution 13, 64–69.

